# Diet-induced changes in metabolism influence immune response and viral shedding dynamics in Jamaican fruit bats

**DOI:** 10.1101/2023.12.01.569121

**Authors:** Caylee Falvo, Dan Crowley, Evelyn Benson, Monica N. Hall, Benjamin Schwarz, Eric Bohrnsen, Manuel Ruiz-Aravena, Madison Hebner, Wenjun Ma, Tony Schountz, Agnieszka Rynda-Apple, Raina K. Plowright

**Affiliations:** Department of Public and Ecosystem Health, College of Veterinary Medicine, Cornell University, Ithaca NY 14853 USA; Department of Microbiology & Cell Biology, Montana State University, Bozeman MT 59715 USA; Research and Technologies Branch, National Institute of Allergy and Infectious Diseases, National Institutes of Health, Hamilton MT 59840 USA; Department of Veterinary Pathobiology, College of Veterinary Medicine and Department of Molecular Microbiology and Immunology, School of Medicine, University of Missouri, Columbia MO 65211 USA; Center for Vector-Borne Infectious Diseases, Department of Microbiology, Immunology, and Pathology, Colorado State University, Fort Collins CO 80523 USA; Department of Wildlife, Fisheries and Aquaculture, Mississippi State University, Starkville, MS 39762 USA (MRA current address)

**Author notes:** Contributed equally (Caylee Falvo, Dan Crowley). Senior authors (Agnieszka Rynda-Apple, Raina K. Plowright).

## Abstract

Land-use change may drive viral spillover from bats into humans, partly through dietary shifts caused by decreased availability of native foods and increased availability of cultivated foods. We manipulated diets of Jamaican fruit bats to investigate whether diet influences shedding of a virus they naturally host. To reflect dietary changes experienced by wild bats during periods of nutritional stress, bats were fed either standard or putative suboptimal diets which were deprived of protein (suboptimal-sugar) and/or supplemented with fat (suboptimal-fat). Upon H18N11 influenza A-virus infection, bats fed the suboptimal-sugar diet shed the most viral RNA for the longest period, but bats fed the suboptimal-fat diet shed the least viral RNA for the shortest period. Unlike mice and humans, bats fed the suboptimal-fat diet displayed higher pre-infection levels of metabolic markers associated with gut health. Diet-driven heterogeneity in viral shedding may influence population-level viral dynamics in wild bats and alter risk of shedding and spillover to humans.

---

Zoonotic spillovers may be occurring more frequently, with anthropogenic land-use change cited as a major driver ^1,2^. Land-use change impacts wildlife habitat structure, resource availability, and animal behavior ^1,3^. Changing these ecological conditions can increase risk of spillover events, both through increasing contact rates between humans and wildlife and by impacting the immune systems of wildlife reservoir hosts^4^. Although the direct mechanisms linking land-use change to viral spillover are multivariate, changes in food availability leading to dietary shifts or nutritional stress are likely to play an important role.

Food availability, dietary shifts, and nutritional stress can impact the immune response. Immune response must compete with other physiological processes when food and energetic resources are limited ^5^. Nutritional stress, such as a caloric deficiency or imbalance, diverts resources from the immune system ^6^ and increases susceptibility to a variety of infections ^7^. Studies in animal models have shown that low protein diets increase likelihood of virus shedding (mice, pigs) ^8,9^, increase disease severity (mice) ^10^, and reduce survival (mice) ^10^. High fat diets can increase infection severity and duration of viral shedding (hamsters) ^11^. Although a link between diet composition, viral infection, and shedding patterns has been established in these animal model systems, no such link has been established in a controlled setting in a natural reservoir of many zoonotic viruses such as bats.

For wild bats, a widespread alteration of habitat causes changes in roosting and foraging behavior, including a switch to anthropogenic food sources ^12–15^. In Australian flying foxes bats, zoonotic spillovers of Hendra virus into horses is driven by habitat loss which leads to acute shortages of nectar and pollen, and bats experience acute nutritional stress ^12^. During these nectar shortages, bats switch from feeding on native eucalyptus nectar and pollen to foods in anthropogenic landscapes including a variety of cultivated plants (e.g., orange, cocos palm, camphor laurel)^15–17^. The nutritional composition of cultivated foods differs substantially from native nectar and pollen. Nectar is mainly composed of sugars (glucose, fructose, sucrose) and eucalypt pollen contains protein, but negligible lipids ^18^. Cultivated plants that bats have been observed eating have lower protein and/or higher lipid content ^19,20^. Variation in dietary protein and lipid content influence viral shedding patterns experimentally in model species ^8,9,11^. Because spillovers of Hendra virus from flying foxes into horses follow these nutritional stress events, it is hypothesized that poor nutrition increases viral shedding rates which leads to spillovers. However, like most studies in wildlife, these findings are correlational, and it is difficult to establish causality without controlled lab-based experiments.

To directly test the effects of dietary changes on viral shedding in bats, we infected Jamaican fruit bats (*Artibeus jamaicensis*; JFBs) with a virus that *Artibeus* bats naturally host, H18N11 influenza A virus (H18N11-IAV). We assigned naïve JFBs to diets designed to mimic dietary changes observed in the Hendra virus system^12^, but which are believed to be representative of nutritional limitations associated with dietary shifts in other bats that are confirmed or implied reservoirs of zoonotic viruses (e.g., *Rousettus aegyptiacus*) ^14^. JFBs were then infected with H18N11-IAV, to directly address whether and how the suboptimal diets would increase viral shedding by inducing changes in metabolism and subsequently the innate immune response. As such, we examined the metabolome of the bats to provide a broader understanding of the influence of diet on their overall physiological status.

## Results

### Interaction between infection and diet influences food consumption

The JFBs’ weights did not change from the beginning of the experiment until immediately pre-infection (experimental timeline Figure 1A) in any of the diet groups (Figure 1B). Post-infection, suboptimal-sugar and standard diet bats gained a small amount of weight (0.06 g/day average), while bats on the suboptimal-fat diet lost a small amount of weight (-0.02 g/day average). Despite the random selection of bats for each diet group, weights varied by treatment and the standard diet bats were slightly heavier on average (ANOVA, DF = 2, F = 5.289, p = 0.013).

**Figure 1.**
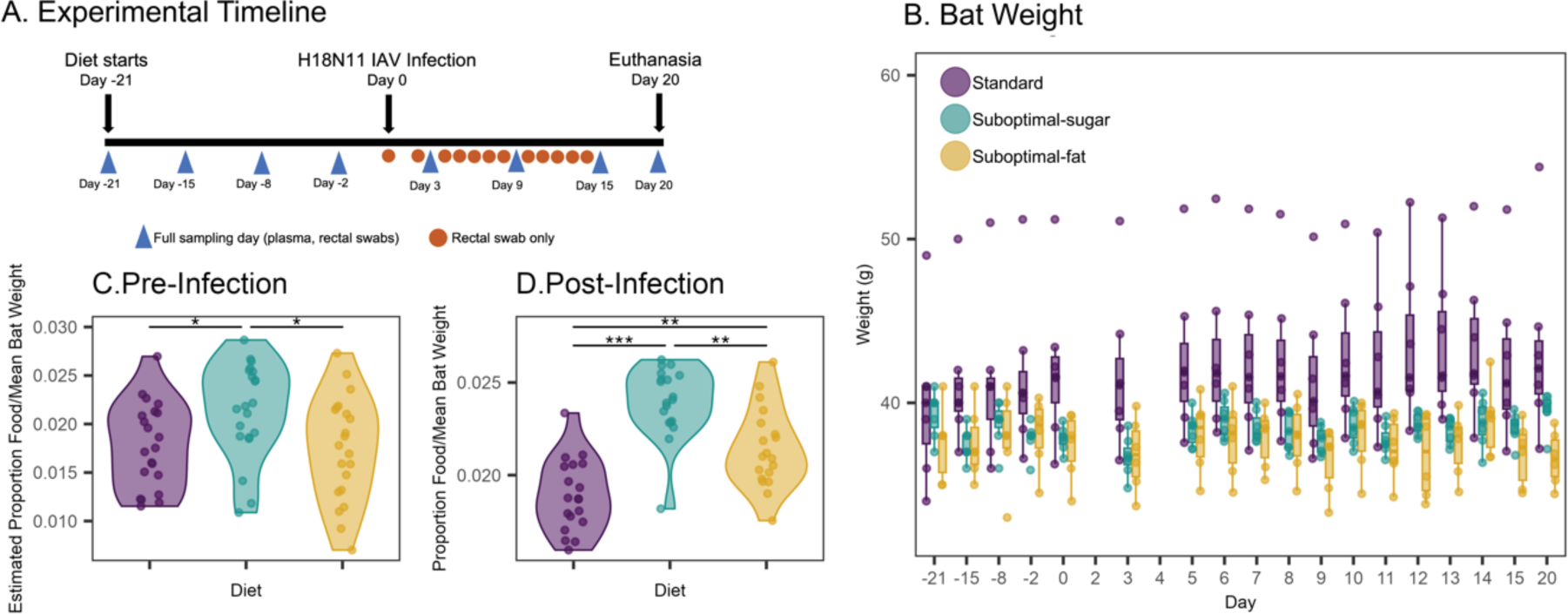
Interaction between infection and diet influences food consumption. (A) Schematic of the timeline of the experiment. (B)Weight of bats throughout the experiment. Standard diet bats were heavier on average, but between day -21 (diet start) and day 0 (infection) weight did not significantly vary. Post-infection (day 0 to day 20), standard and fruit diet bats gained a small amount of weight (lme, 0.06 g/day estimate; df = 257, t = - 3.92, p = 0.001) while suboptimal-fat diet bats lost a small amount of weight (lme, -0.02 g/day; df = 257, t = -3.39, p = 0.008). (C) Pre-infection overall differences in estimated proportion of food consumed, based on using the average food provided minus weight of food leftover, corrected by average weight of bats in the cage. Suboptimal-sugar diet cage consumed more food than both the suboptimal-fat and standard cages (ANOVA, F = 4.32, df = 2, p = 0.02). (D) Post-infection overall food consumed, based on actual weight of food provided minus weight of food leftover, corrected by average weight of bats in the cage. Both suboptimal diet cages consumed more food overall than standard diet bats, although the suboptimal-sugar bats still consumed the most (ANOVA, F = 30.36, df = 2, p < 0.001). *Boxplots show the median, 25^th^ and 75^th^ percentiles, and the range or 1.5 * IQR.* *Asterisks indicate:* p < 0.0005 = ***, p < 0.005 = **, p < 0.05 = *, p < 0.1 =.

We measured food consumption to determine changes in appetite. Pre-infection, the suboptimal-sugar diet cage consumed the most food (Figure 1C). Post-infection, both suboptimal diet cages consumed more food overall than standard diet bats, although the suboptimal-sugar bats still consumed the most (Figure 1D). Normalizing by number of bats per cage did not substantially change results (Supp. Figure 1). Aside from the change in food consumption, the infection did not result in other clinical symptoms in bats.

### Diet led to unique metabolomic profiles in rectal environment, plasma, and liver

We analyzed rectal swabs, plasma, and liver samples of individual bats to determine how diet affected systemic and local metabolic profiles. We found that manipulating diet led to distinct metabolic profiles pre-infection in both the rectal environment and plasma (sPLSDA; Figure 2A, 2C). These metabolic groupings appeared stable by the point of infection and were largely maintained between diets post-infection (Supp. Figure 2).

**Figure 2.**
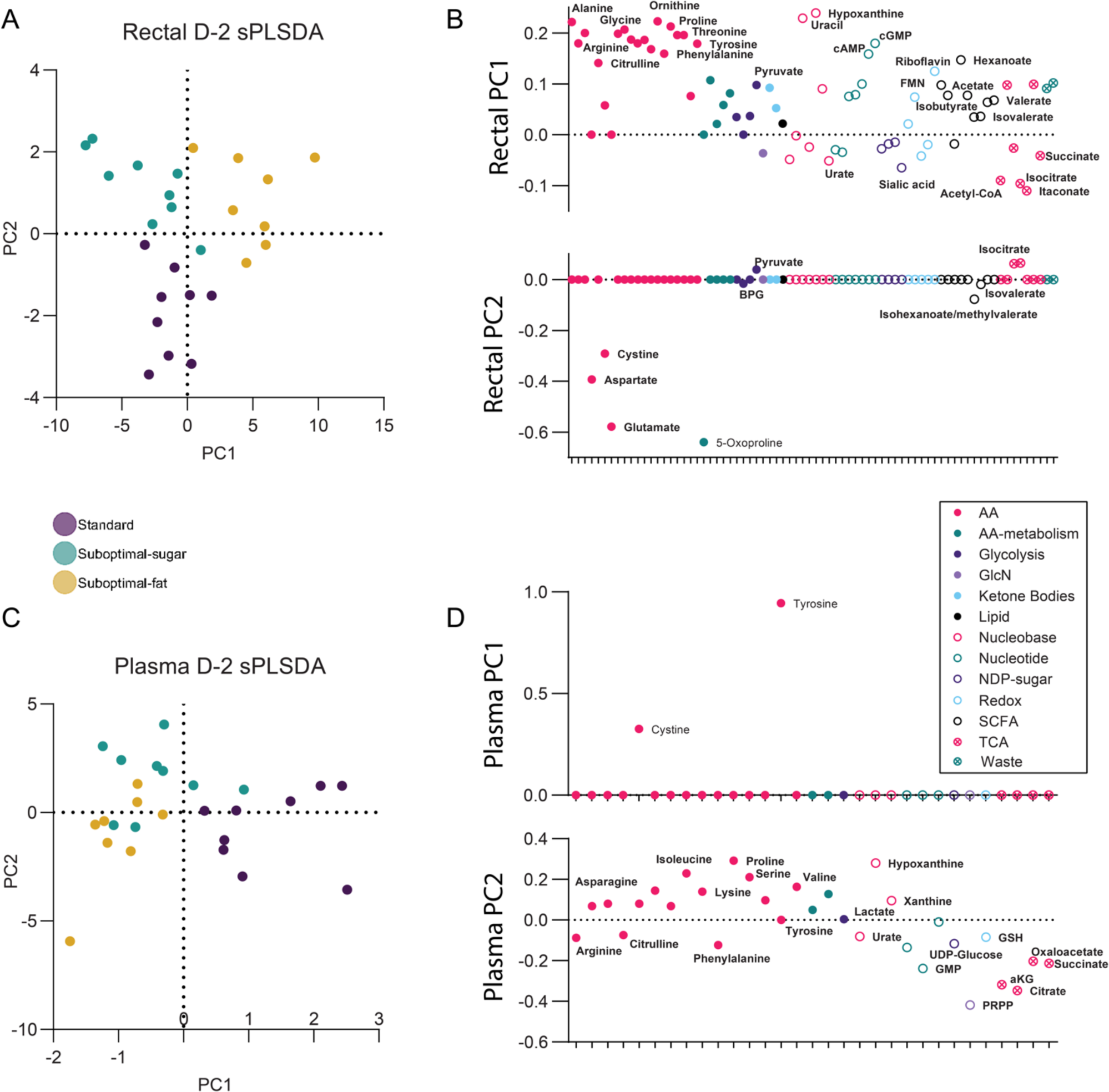
Diet led to unique metabolomic profiles in rectal environment and plasma. (A, C) Sparse partial least squares regression (sPLSDA) classification of bats in each diet pre-infection (day -2) based on metabolites in rectal swabs (A) and plasma (C). Each point represents an individual bat. (B,D) Loadings of metabolites from first two axes of the sPLSDA (PC1 and PC2) for rectal swabs (B) and plasma (D). Each point represents a metabolite.

In the rectal swabs pre-infection (day -2), the suboptimal-fat diet was distinct from the other diets based on PC1 (Rectal PC1, Figure 2B), whereas both suboptimal diets were distinguishable from the standard diet based on PC2 (Rectal PC2, Figure 2B). Divergence between diets was partly driven by differences in metabolites known to correlate with gut health and immune function including short chain fatty acids (SCFAs), amino acids, TCA (tricarboxylic acid cycle) metabolites, basic sugars, and nucleobases. Suboptimal-fat diet bats had decreased levels of several TCA-related metabolites (e.g., acetyl-CoA, succinate, isocitrate, and cis-aconitate), suggesting different prioritization of energy metabolism in the host and/or microbial compartment (Rectal PC1, Figure 2B). Suboptimal-fat diet bats also had higher levels of most amino acids, SCFAs, and nucleobases/nucleosides (e.g., cGMP, xanthine, hypoxanthine), suggesting increased production or turnover of protein and nucleobases/nucleosides. Both suboptimal diets had decreased levels of 5-oxoproline, glutamate, aspartate, and cystine (Rectal PC2, Figure 2B; Supp. Figure 2A). Surprisingly, the availability of most amino acids (except glutamate, aspartate, and cystine) were highest in suboptimal-fat diet bats, but lowest in suboptimal-carbohydrate diet bats, despite both suboptimal diets being protein poor (Supp. Figure 3A).

In the plasma, the suboptimal diets clustered together but were distinguishable from the standard diet (Plasma PC1, Figure 2C). The suboptimal diets had lower levels of several amino acids compared to the standard diet. This pattern contrasts with the elevation of amino acids in the rectal environment of suboptimal-fat diet bats. Several nucleobases (hypoxanthine, xanthine) were lowest in the plasma of suboptimal-fat diet bats (Supp. Figure 3B) despite being elevated in their rectal environment (Supp. Figure 3A). The opposing patterns of amino acids and hypoxanthine between the rectal and plasma metabolome in the suboptimal-fat diet, but not in the suboptimal-sugar bats, suggests a mechanism in suboptimal-fat diet bats with regard to the transport, synthesis, or recycling of these base nutrients between host and microbes in the gut compartment. The suboptimal diets were not as clearly distinguished from each other in the plasma (Figure 2C), except for an elevation of citrulline and arginine in the suboptimal-fat bats relative to the suboptimal-sugar bats (Plasma PC2, Figure 2D).

Diet-dependent patterns were easily discernible in the liver and metabolic profiles reflected the main source of energy the bats were receiving (Supp Figure 3A, 3B). Like plasma, both suboptimal diets were amino acid depleted in the liver as expected from a protein-poor diet (Supp. Figure 3C). The notable exception was arginine, which was elevated in both suboptimal diets compared to the standard diet bats. Patterns consistent with higher sugar-based metabolism were apparent in both suboptimal diets, which was expected for the suboptimal-sugar diet but surprising for the suboptimal-fat diet. However, this pattern could also indicate higher levels of gluconeogenesis in the suboptimal-fat bats (as opposed to glycolysis), as directionality cannot be determined for bidirectional pathways via these data. In line with expectations, levels of acetyl-CoA were highest in the suboptimal-fat diet bats which suggests elevated fatty acid oxidation. Higher levels of the ketone bodies 3-hydroxybutyrate and acetoacetate in the standard diet compared to the suboptimal diets suggests bats may use ketogenic metabolism to a higher degree when consuming a protein-supplemented diet.

**Figure 3.**
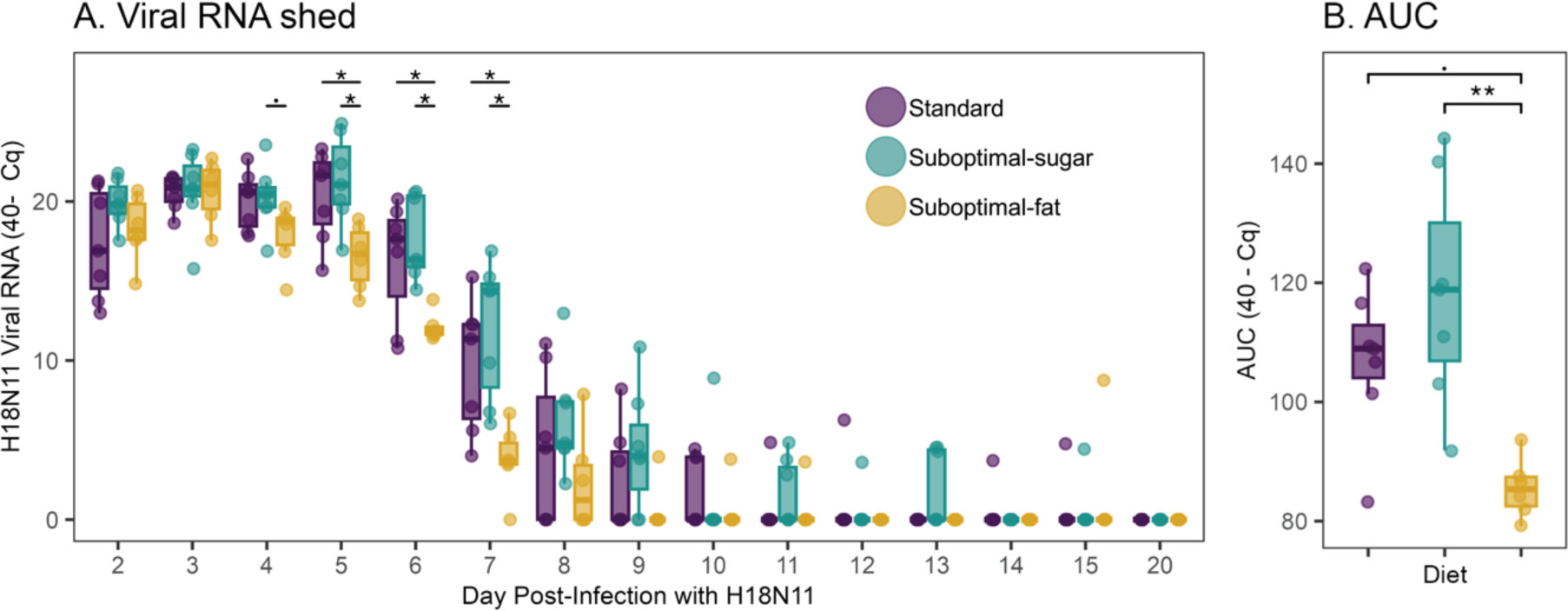
Diet influences the duration and quantity of H18N11 viral RNA shedding. (A) Viral RNA detected post-infection (inverse Cq). Differences in viral RNA were different among diets on days 4-7 (ANOVA; day 4 p = 0.08, F = 2.911; day 5 p = 0.0107, F = 6.003; day 6 p = 0.00619, F = 6.959; day 7 p = 0.00362, F = 2.7.968). (B) Area under the curve (AUC) which sums the total amount of viral RNA shed varied significantly by diet (ANOVA p = 0.004, F = 7.628). Suboptimal-fat diet bats cumulatively shed less than both suboptimal-sugar (Tukey HSD, p = 0.003) and standard diet bats (Tukey HSD, p = 0.054). *Boxplots show the median, 25^th^ and 75^th^ percentiles, and the range or 1.5 * IQR.* *Asterisks indicate:* p < 0.0005 = ***, p < 0.005 = **, p < 0.05 = *, p < 0.1 =.

### Diet influences duration and quantity of H18N11 viral RNA shedding

Next, we determined whether there were differences in H18N11-IAV viral RNA (vRNA) shedding in the diet groups. There were no significant differences between the standard and suboptimal-diet bats on any day, though suboptimal-sugar diet bats trended toward higher levels of vRNA shedding on all days measured (Figure 3A). Unexpectedly, the bats on the suboptimal-fat diet shed less vRNA on days 4 - 7. While not significant, the average duration of shedding between diets followed the same trend (suboptimal-sugar = 8.3 days, standard = 7.7, suboptimal-fat = 6.7). Bats continued to shed through day 15, although no individual bat shed continuously.

Using area under the curve (AUC) analysis to quantify total vRNA shed per bat, we found that total shedding varied significantly by diet (Figure 3B). Standard diet bats cumulatively trended toward shedding less vRNA than suboptimal-sugar diet bats, but the difference was not significant. Suboptimal-fat diet bats cumulatively shed less vRNA than both suboptimal-sugar and standard diet bats.

### Gross Tissue Examination

We next examined whether and how diet affected appearance of lymphoid tissue post-infection (Table 1). None of the bats on the standard or suboptimal-fat diet had visible Peyer’s patches, but two in the suboptimal-sugar diet group had large, distinct Peyer’s patches. All infected bats in the standard diet group had visible mesenteric lymph nodes (MLNs). Only one suboptimal-sugar diet and three suboptimal-fat diet bats had visible MLNs.

**Table 1.** Gross lymphoid tissue counts. Counts of Peyer’s patches and mesenteric lymph nodes (MLNs) found during necropsy examination in each bat, including infected bats and uninfected control bats.

|  | Peyer's patches | Mesenteric lymph nodes |  |
| --- | --- | --- | --- |
| Standard | 0/7<br>0/2 | 7/7<br>0/2 | infected bats<br>control bats |
| Suboptimal-sugar | 2/7<br>0/2 | 1/7<br>0/2 | infected bats<br>control bats |
| Suboptimal-fat | 0/6<br>0/2 | 3/6<br>1/2 | infected bats<br>control bats |

### Diet but not infection influences rectal TNF expression

Because we found that diet affected the quantity and duration of vRNA shedding, we next tested whether infection affected the antiviral and/or inflammatory responses.

Pre-infection (day 0), TNF expression was highest in the standard diet bats and lowest in the suboptimal-fat diet bats (Figure 4A), suggesting that diet influenced inflammatory state, regardless of infection. Although TNF expression in the standard diet bats was slightly higher than the suboptimal diets on day 3, there were no other within-day differences in TNF expression post-infection. Levels of Tgfb and Ifng were slightly higher in the standard diet bats relative to the suboptimal diets on several days (Figure 4B, 4C; day 2, 6 Tgfb; day 3 Ifng).

**Figure 4.**
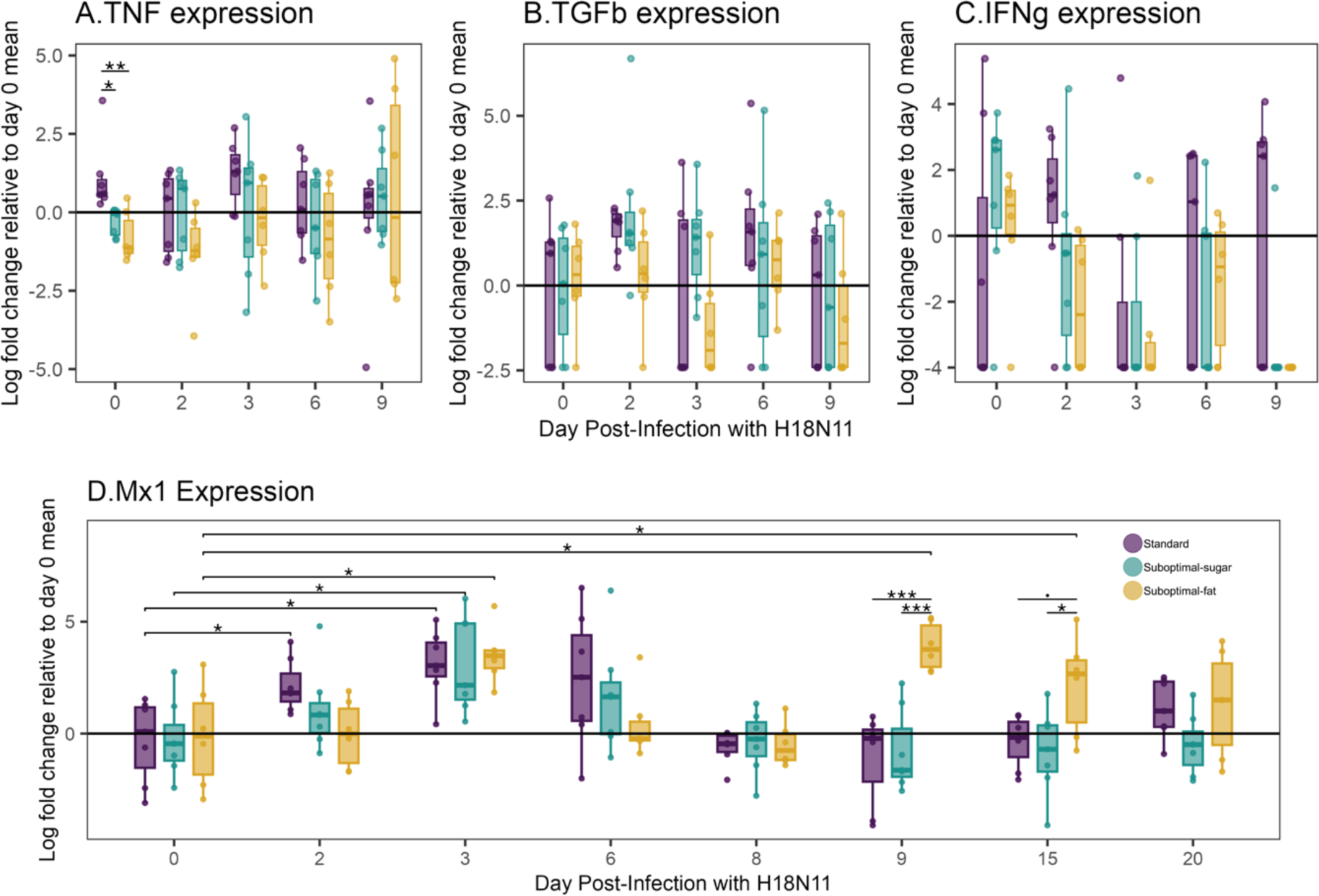
Diet but not infection influence TNF expression, while antiviral protein Mx1 has diet-specific expression pattern. Gene expression from rectal swabs. Values are the natural log-fold change in gene expression relative to day 0 mean expression. (A) *TNF* expression on days 0, 2, 3, 6, and 9. On day 0 there were significant differences in expression between bats on different diets (ANOVA, F = 8.692, corrected p-value = 0.03). (B) *TGFb* expression on days 0, 2, 3, 6, and 9. (C) *IFNg* expression on days 0, 2, 3, 6, and 9. (D) *Mx1* expression on days 0, 2, 3, 6, 8, 9, 15, and 20. There were within-diet differences on day 9 (ANOVA, F = 16.17, t = 2, corrected p-value = 0.0008) and day 15 (ANOVA, F = 4.99, t = 2, corrected p-value = 0.069). Compared to their day 0 baseline level, standard diet bats increased expression of Mx1 on day 2 and day 3 (paired t-test, day 2 t = -2.85, df = 6, corrected p-value = 0.049; day 3 t = -4.17, df = 6, corrected p-value = 0.029). Suboptimal-sugar diet bats also increased expression on day 2 and day 3 (day 2 paired t-test; t = -2.17, df = 6, corrected p-value = 0.10; day 3 t = -4.33, df = 6, corrected p-value = 0.03). However, suboptimal-fat diet bats had a delayed response and did not increase Mx1 expression until day 3 (paired t-test: t = -3.27, df = 5, corrected p-value = 0.04). For all diets, Mx1 expression peaked at day 3 and decreased back to baseline at day 8. However, in suboptimal-fat diet bats, a second peak of Mx1 occurred at day 9 (paired t-test, t = -3.46, df = 5, corrected p-value = 0.04) and did not return to baseline levels until day 20 (day 15 paired t-test, t = -3.45, df = 5, corrected p-value = 0.04). *Boxplots show the median, 25^th^ and 75^th^ percentiles, and the range or 1.5 * IQR.* *Asterisks indicate p-values:* p < 0.0005 = ***, p < 0.005 = **, p < 0.05 = *, p < 0.1 =.

We determined whether cytokine expression was influenced by infection by comparing expression relative to the day 0 baseline. On day 2 post-infection, there were slight changes in expression of Tgfb and Ifng in most bats. Standard diet and suboptimal-sugar bats had small increases in Tgfb (Figure 4B). Standard diet bats had a small increase in Ifng while both the suboptimal diets had a small decrease in Ifng (Figure 4C). On days 3-9, most bats had undetectable levels of Ifng. Overall, it appears that diet led to differences in baseline expression, but H18N11-IAV infection induced only moderate responses in the cytokines measured.

### Antiviral protein Mx1 had diet-specific expression patterns

Mx1 is an interferon-stimulated antiviral protein which is important for an efficient anti-IAV immune response in humans and mice ^21^. Pre-infection Mx1 expression was similar between the bats fed different diets, although all bats were expressing detectable Mx1 (Figure 4D).

Relative to baseline levels (day 0), Mx1 expression increased more rapidly in the standard and suboptimal-sugar diet bats on days 2 and 3. Suboptimal-fat diet bats had a delayed response and did not increase Mx1 expression until day 3. For all diets, Mx1 expression peaked at day 3 and decreased back to baseline at day 8. However, in suboptimal-fat diet bats only, a second peak of Mx1 occurred at day 9 and did not return to baseline levels until day 20 (Figure 4D).

### Infection-driven metabolic changes are diet-specific in the gut but not in circulation

We next looked for correlations between the metabolites and viral shedding. We began by determining which metabolic changes were triggered by viral infection in the gut (rectal swabs; Figure 5A) or in circulation (plasma; Figure 5B).

**Figure 5.**
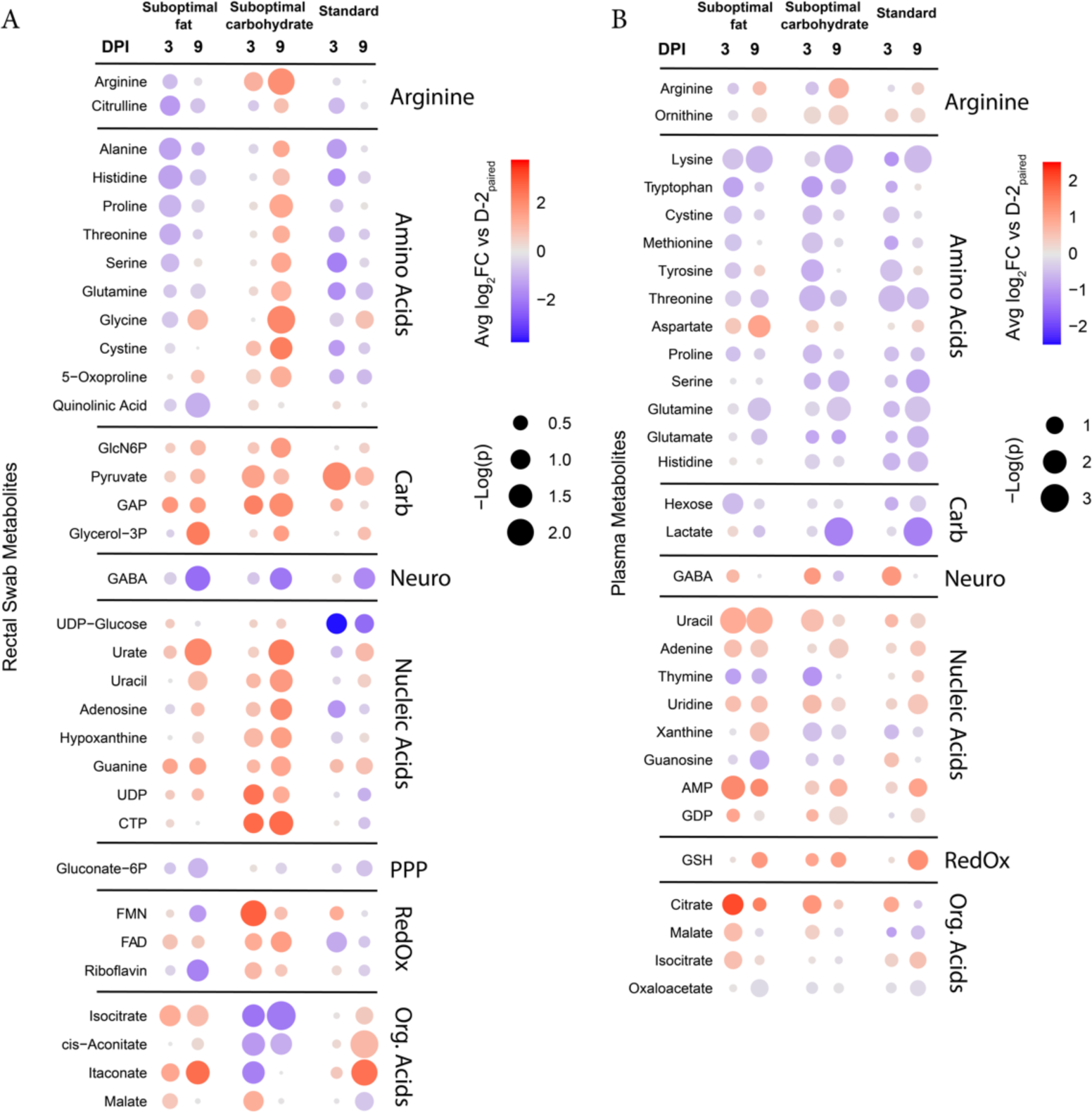
Infection-driven metabolic changes are diet-specific in the gut but not in circulation. Metabolites in the (A, left) rectal environment and (B, right) plasma that changed relative to pre-infection (day-2) levels on days 3 and 9 post-infection (when bats were still shedding detectable H18N11 vRNA). Color indicates log2-fold change relative to day-2 values and size of dot indicates -log(p value). This analysis was performed separately for each diet group due to the differences in the metabolite levels pre-infection. Infected bats were also compared to uninfected bats on each day post-infection to adjust for metabolic changes over the span of the experiment.

The rectal compartment displayed infection-triggered responses in amino acids, carbohydrate metabolism, nucleic acids, flavin-associated cofactors, and TCA-associated organic acids (Figure 5A). Few of these responses were conserved across diets, except for the changes in carbohydrate metabolism. The suboptimal-fat and standard diet bats had a decrease in amino acids on day 3. The suboptimal-sugar diet bats had an opposite pattern, with a strong increase in amino acids by day 9. Suboptimal-sugar diet bats also displayed a more robust increase in nucleic acid and nucleic acid metabolites throughout infection compared to the other diets. An exception was urate, which showed increases in both suboptimal diets at day 9 post-infection.

In contrast to the gut, post-infection metabolic patterns in the plasma were largely conserved between diets, but included changes in amino acids, carbohydrate metabolism, nucleic acids, and organic acids (Figure 5B). While the magnitude differed between diets, the directionality and timing were largely preserved.

### Gut levels of amino acids pre- and post-infection are negatively correlated with viral shedding

We next asked which pre-infection metabolite levels and infection-associated metabolic changes had explanatory potential for the differences observed in viral shedding between diets. To this end, we looked for correlations of individual viral AUC with metabolite profiles on day -2 and each day following for which metabolite data was available and vRNA was detectable (day 3 and 9; Figure 6A, 6B). Diet was not considered in this calculation and each infected bat was treated individually.

**Figure 6.**
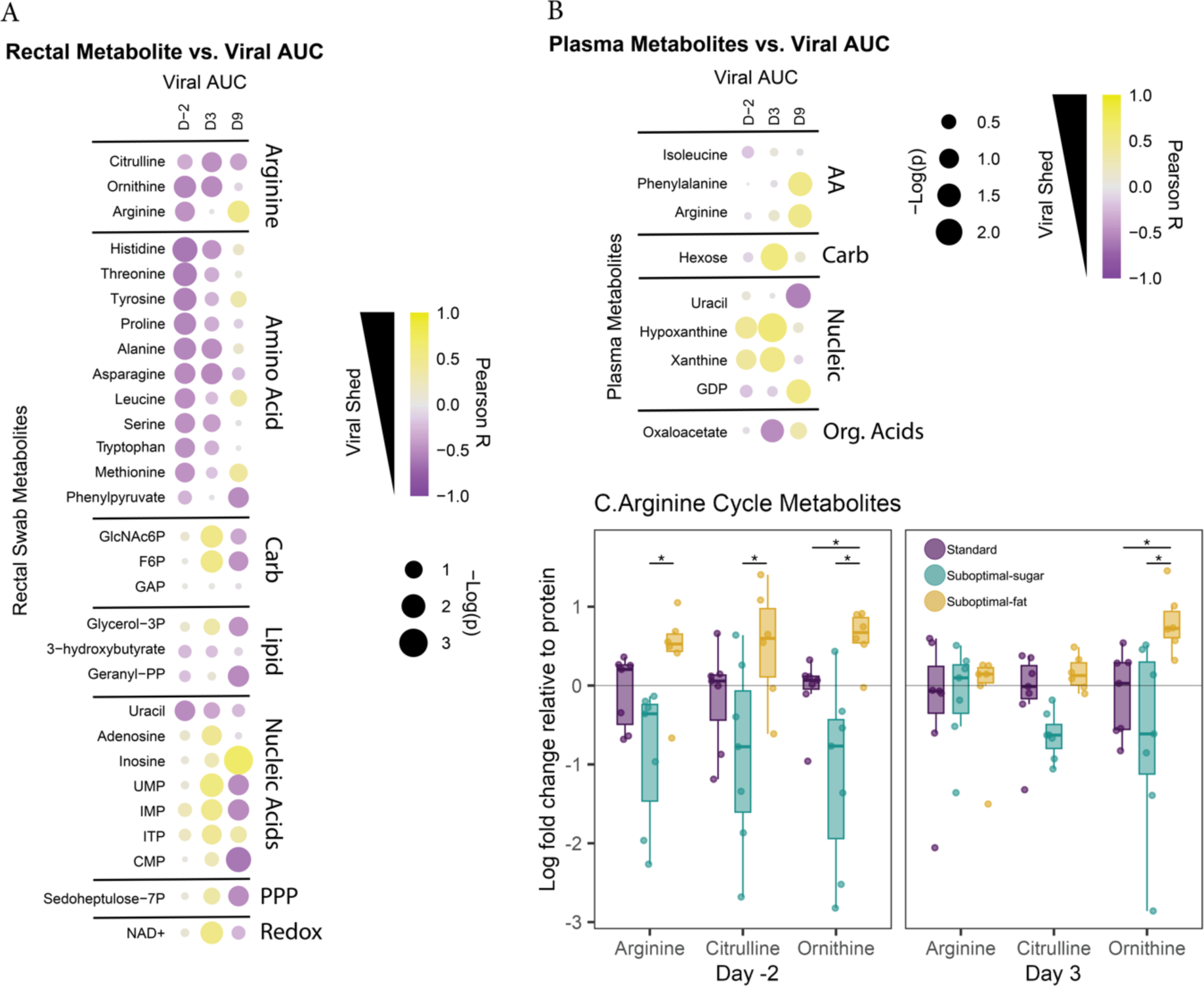
Amino acid levels in the gut pre- and post-infection are negatively correlated with viral shedding. Metabolites from the (A) rectal environment and (B) plasma that are correlated with viral AUC for each bat on day -2 (pre-infection) and each day following for which metabolite data was available while virus was still detectable (day 3 and day 9). Diet was not considered in this calculation and instead each infected bat was treated individually. Color indicates positive correlation with AUC (i.e., higher levels of metabolite correlated with higher total virus shed) versus negative correlation with AUC. Size of dots indicates -log(p-value). (C) Levels of citrulline, arginine, and ornithine pre-infection (day -2) and post-infection (day 3) from the rectal environment. Data are standardized to the protein mean of that day to account for batch effects. *Boxplots show the median, 25^th^ and 75^th^ percentiles, and the range or 1.5 * IQR.* *Asterisks indicate:* p < 0.0005 = ***, p < 0.005 = **, p < 0.05 = *, p < 0.1 =.

Pre-infection levels of amino acids in the gut were strongly negatively correlated with viral shedding AUC (Figure 6A), consistent with the higher rectal levels of amino acids in the lower shedding suboptimal-fat diet bats. This correlation continued out to day 3 post-infection, near the peak of vRNA shedding, and was lost by day 9, consistent with the infection-driven increase in the amino acids in the suboptimal-sugar diet bats at day 9 (Figure 6A). While no pre-infection gut metabolites were positively correlated with viral AUC, day 3 carbohydrate metabolites and nucleic acids positively correlated with viral AUC.

Of particular interest, we found a strong negative correlation pre-infection of viral AUC with all three members of the arginine cycle (arginine, citrulline, and ornithine). Pre-infection, levels of all three metabolites were higher in the suboptimal-fat diet bats (Figure 6C), suggesting that diet influenced the arginine cycle.

Plasma metabolites were less correlated with viral AUC, consistent with the observed conservation of infection-triggered plasma metabolite patterns between diets (Figure 6B). Notably, plasma hypoxanthine and xanthine levels both pre-infection and on day 3 were positively correlated with viral AUC, consistent with these metabolites being lowest in the plasma of suboptimal-fat diet bats.

### Suboptimal-fat diet bats have distinct plasma antioxidant response

Because we determined that diet led to systemic changes in metabolites associated with redox processes (e.g., xanthine, hypoxanthine, flavin cofactors), we measured enzymatic activity of glutathione peroxidase (GPx) and superoxide dismutase (SOD) in hemolysate samples.

Pre-infection, Gpx and SOD activity was similar in all diets (Figure 7). However, on day 3 post-infection, the suboptimal-fat diet bats had the lowest levels of GPx (Figure 7A) and the highest levels of SOD (Figure 7B). While GPx activity returned to baseline by day 9 in suboptimal-fat diet bats, SOD remained elevated. Despite metabolic signatures of oxidative stress in the rectal metabolome of suboptimal-sugar diet bats and in the plasma metabolome of standard diet bats (i.e., hypoxanthine, xanthine), we observed no differences in these diets.

**Figure 7.**
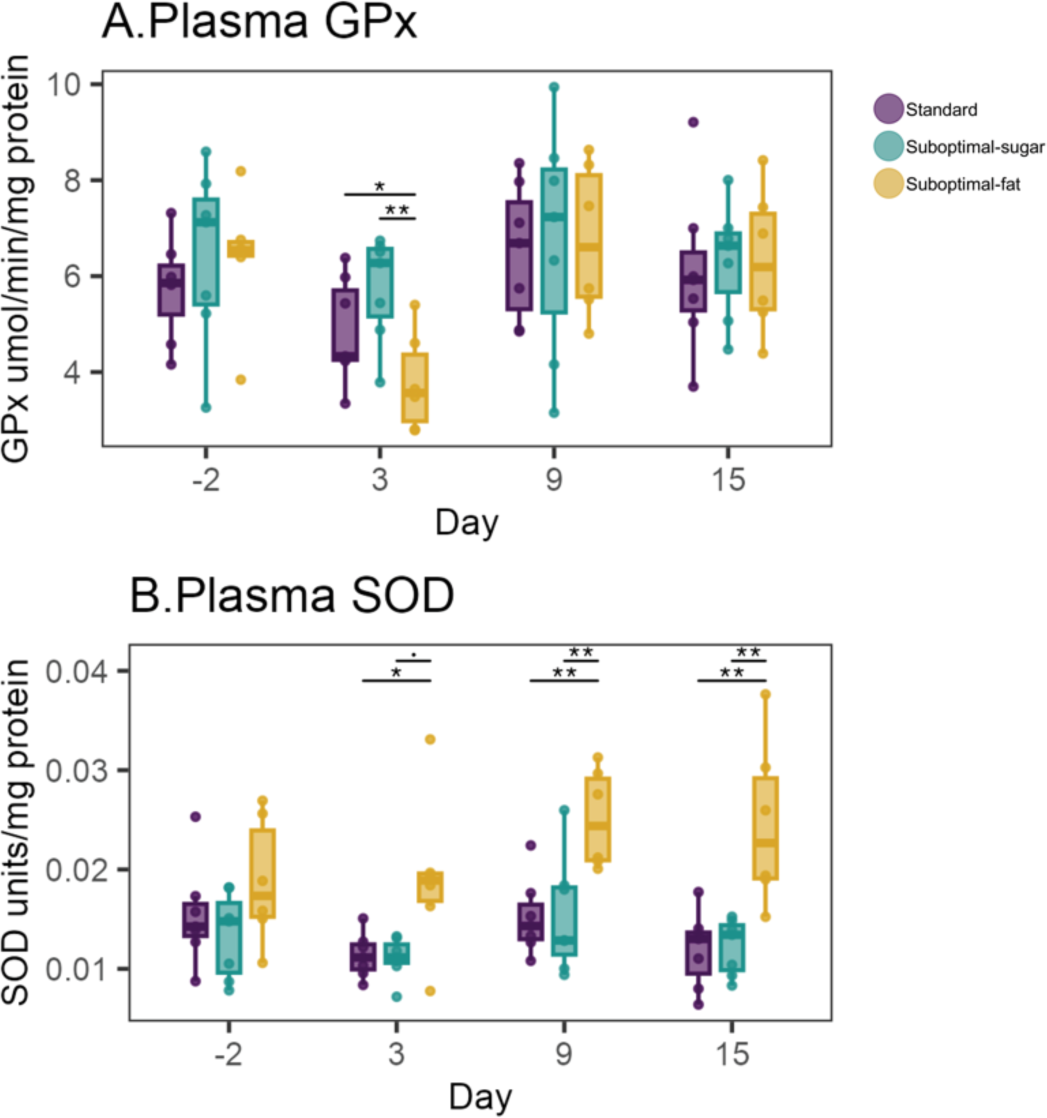
Suboptimal-fat diet hats have distinct plasma antioxidant response. Plasma levels of antioxidant enzymes pre-infection (day -2) and post-infection on days 3, 9, and 15. (A) Glutathione peroxidase (GPx) activity pre-infection (day-2) was similar across diets (F = 0.5959, corrected p-value = 0.99). On day 3 post-infection, the suboptimal-fat diet bats had the lowest levels of GPx (F =8.70, corrected p-value = 0.006). GPx activity standardized to total protein in each sample. (B) Superoxide dismutase (SOD) activity pre-infection (day-2) was similar across diets (F = 1.463, corrected p-value = 0.2525). On day 3 post-infection, the suboptimal-fat diet bats had the highest levels of SOD (Figure 7B; F = 4.12, corrected p-value = 0.04; data were log transformed for statistical test). SOD activity standardized to total protein in each sample. *Boxplots show the median, 25^th^ and 75^th^ percentiles, and the range or 1.5 * IQR.* *Asterisks indicate:* p < 0.0005 = ***, p < 0.005 = **, p < 0.05 = *, p < 0.1 =.

### Discussion

Bats are reservoirs of many viruses that are important to public health, but little is known about the immunological and metabolic drivers of viral shedding. Bats are continuously facing changing landscapes, including loss of native habitat, increasingly forcing them to rely on novel food sources. In this study, we attempted to recapitulate these dietary changes in an experimental setting. As hypothesized, Jamaican fruit bats on the suboptimal-sugar (fruit-only) diet shed more H18N11-IAV viral RNA (vRNA). However, contrary to our expectation, bats on the suboptimal-fat diet shed the least amount of vRNA. We also found that bats on the suboptimal diets consumed more food, despite only small changes in weight. Furthermore, the suboptimal-fat diet bats had distinct patterns of cytokine, antiviral protein, metabolite, and antioxidant enzymes that suggest diet mediated a change in their metabolism and immune system which influenced the pattern of vRNA shedding. The duration of pathogen shedding, or infectious period, is import for understanding how pathogens spread and are maintained in a population ^22^. Our results suggest that periods of dietary stress may alter patterns of pathogen shedding within bat populations, giving support to the hypothesis that diet may be mediating the interaction between land-use change and spillover events.

We observed potential sickness behaviors in our bats, whose appetites were altered by infection. Specifically, bats on the suboptimal-fat diet ate more food than bats on the standard diet, although their weights did not vary substantially. We also observed that the suboptimal-sugar diet bats tended to consume more food than the other diets throughout the experiment, although the difference was greater post-infection. This result was not surprising, as previous studies have shown that JFBs given low-protein diets consume more food than those given high-protein diets ^23^. By weight, the diets differed calorically and nutritionally, so the suboptimal-sugar diet bats may have consumed more to reach the same nutrient or calorie demands. If these trends are similar in wild bats, bats on suboptimal diets may spend more time foraging than bats consuming typical native foods. A previous study of bat movement indeed found that during food shortages, bats increased the number of foraging stops ^15^. The risk of viral spillover increases with contact between hosts ^24^, and more time foraging increases the chance of contact with a secondary host. Future studies are needed to determine whether diet and viral infection change foraging behavior. Nonetheless, our results suggest that in addition to the heterogeneity in viral shedding, there may also be a behavioral change, which further affects viral transmission dynamics within populations ^22^.

We identified metabolic markers of diet and viral shedding that relate to gut health, including citrulline/arginine, several SCFAs, and hypoxanthine. The metabolite patterns in the gut collectively suggest that the suboptimal-fat diet induced a distinct gut metabolic environment pre-infection that may have primed bats’ immune response, leading to less vRNA shedding and a shorter shedding duration. In mice and humans, higher levels of citrulline/arginine, hypoxanthine, and SCFAs are all related to gut health, including increased intestinal barrier integrity ^25,26^, reduced inflammation ^25^, increased healing capacity ^26^, and improved anti-viral immune responses ^27,28^. The effects of these metabolites on local and systemic immune and physiologic function provide a cohesive diet-induced difference in the gut environment, which may be responsible for the differences in viral shedding, although the exact mechanism remains unknown.

Suboptimal-fat diet bats had distinct patterns of cytokine expression, antiviral protein expression, and plasma antioxidant enzyme activity. Pre-infection, rectal levels of TNF were lowest in the suboptimal-fat diet bats, suggesting variation in gut inflammation related to diet. This TNF expression pattern was consistent with the metabolite signature of reduced inflammation in the suboptimal-fat diet bats. TNF is a potent pro-inflammatory cytokine that can be produced by innate cells early during viral infections, but we did not detect changes in TNF in response to infection, consistent with other studies in bats ^29^. In contrast, we observed a clear induction of the antiviral protein Mx1 in response to infection. The standard and suboptimal-sugar diet bats began expressing Mx1 earlier. Additionally, the suboptimal-fat diet bats had a second peak of Mx1. Levels of Mx1 in the suboptimal-fat diet bats remained elevated throughout the experiment, while the other diets returned to their pre-infection baselines. It is unclear what caused the second peak in Mx1, which is generally only expressed in response to type I or III interferons as part of the canonical interferon response ^21,30^. Although we did not detect vRNA at these time points, it is possible there were residual pathogen-associated antigens (i.e., PAMPs) that elicited additional expression of Mx1. In addition to their distinct Mx1 response, plasma SOD activity increased in the suboptimal-fat diet bats in response to infection, which could increase their capacity to prevent the formation of reactive species, suggesting a role of systemic redox balance. Overall, the suboptimal-fat diet bats had several metabolic and immune differences that may have led to the differences in viral shedding.

Data on the metabolic response to a high fat diet (HFD) in humans and mice suggests that this response in bats could be unusual. A HFD in mice leads to decreased intestinal integrity ^31–33^, decreased production of SCFAs ^34^, and increased inflammatory cytokine production ^33^. Here, SCFAs in the suboptimal-fat diet JFBs were increased relative to the other diets pre-infection (Supp. Figure 2A). Although we did not directly measure intestinal integrity, SCFAs ^25^, hypoxanthine ^26^, and citrulline ^27^ were also higher in the suboptimal-fat diet bats and are each associated with increased intestinal integrity. This suggests that while a HFD may induce some common metabolic pathways in mice/humans and bats, there is clearly a distinct mechanism occurring in the intestine of suboptimal-fat diet bats. Differences in the size and number of Peyer’s patches and MLNs between diets further suggest that diet profoundly shifted the immune environment. Future studies could determine whether the effect of diet on viral shedding is unique to gastrointestinal viruses.

This study provides experimental support for a hypothesis that we generated based on observations of wild bats in the zoonotic Hendra virus system, confirming that diet can influence viral shedding patterns, likely through modification of the bats’ metabolic and immune state prior to infection. Our results demonstrate the plausibility of the hypothesis that changes in diet due to habitat degradation are a driver of viral shedding in bats. Although it is not understood how viruses are maintained over time within bat populations, heterogeneity in viral shedding has the potential to prolong outbreaks and contribute to temporal maintenance of pathogens ^22^. Our observation that bats on both suboptimal diets consumed more food post-infection could translate to increased foraging time in wild bats, increasing the risk of contact with secondary hosts. In wild bats, dietary shifts driven by habitat change may therefore modulate the dynamics of zoonotic pathogens and the risk of spillover to humans. Our findings that dietary changes can both influence the duration of viral shedding, and potentially impact foraging time, have implications for how viral transmission dynamics among wild bats are modeled and understood, and highlight the need to consider ecological factors when studying bat-borne diseases.

## Acknowledgements

We thank Julia R. Port and Catherine M. Bosio for their help on this project. This work was funded by the U.S. National Science Foundation (EF-2133763/EF-2231624), the DARPA PREEMPT program Cooperative Agreement (D18AC00031; DEB-1716698) and NIH (1R01AI134768). The content of the information does not necessarily reflect the position or the policy of the U.S. government, and no official endorsement should be inferred.

## Competing Interests

The authors declare no competing interests.

## Online Methods

### Bat Husbandry & Diet

Male Jamaican fruit bats (*Artibeus jamaicensis*; JFBs) were obtained from a specific pathogen-free breeding colony at Colorado State University and acclimated for three weeks. All care and procedures were in accordance with NIH, USDA, and the Guide for the Care and Use of Laboratory Animals (National Research Council, 2011). Animal protocols were reviewed and approved by the MSU Institutional Animal Care and Use Committee (IACUC 2021-183-IA). MSU is accredited by the Association for Assessment and Accreditation of Laboratory Animal Care (AALAC; accreditation no. 713).

Bats were housed on a 12 hour on/off light dark cycle at 75 +/- 2 degrees F (approx. 23.8 degrees C) and 30-70% humidity (MSU IACUC 2021-183). Bats were randomly assigned to diet groups. Throughout the experiment, bats in each cage ate a distinct diet: a standard diet (fruit with protein supplement: Mazuri Exotic Animal Nutrition SKU 0053414), a suboptimal-sugar diet (fruit alone without any supplement), or a suboptimal-fat diet (fruit with a fat supplement: coconut oil). Fruit given to each cage was identical except for the supplement, and bats were maintained on this diet for 21 days prior to infection (d-21 to d0) and 20 days post-infection (d0 to d20). Bats were randomly assigned to diet groups and to infected versus control groups.

To monitor consumption, leftover food was weighed to determine if there were notable differences in food consumed at a cage level. After infection with H18N11, food was weighed both before and after it was given to the bats every day to more precisely monitor changes in consumption. Control bats were housed separately from infected bats post-infection. Bats were monitored for 20 days post-infection and then euthanized.

### H18N11 Infection

The A/flat-faced bat/Peru/033/2010 (H18N11) was rescued by using eight plasmid reverse genetic system as described previously ^35,36^. The rescued H18N11 virus was amplified in RIE1495 cells and titrated in MDCK cells^35^. After 21 days on their respective diets, we infected the JFBS with the H18N11 virus. We inoculated each JFB with 5 x 10^^5^ TCID_50_ of the H18N11 virus. We selected two JFBs from each diet group to act as controls, inoculating them with an equal volume of sterile saline.

### RNA qRT-PCR

#### Rectal swabs: gene expression & viral RNA

Rectal swabs (Puritan 6” Sterile Mini-tip swabs #25-800-1PD-50) were submerged in DNA/RNA Shield (Zymo R1100) immediately after collection. Nucleic acids were extracted from rectal swabs using Zymo Quick-DNA/RNA Pathogen Miniprep kit following manufacturer’s instructions. Viral RNA was measured using TaqMan Fast Virus 1-Step RT-PCR standard parameters. Primers were designed for the NP region (per Ciminski et al 2019). Cytokine expression levels were measured using iTaq Universal SYBR Green One-Step Kit (BioRad #172-5151) after being treated with DNAse (Turbo DNA-Free Kit, Ambion #AM1907) using standard PCR conditions. [check w evelyn or write out all the PCR conditions]

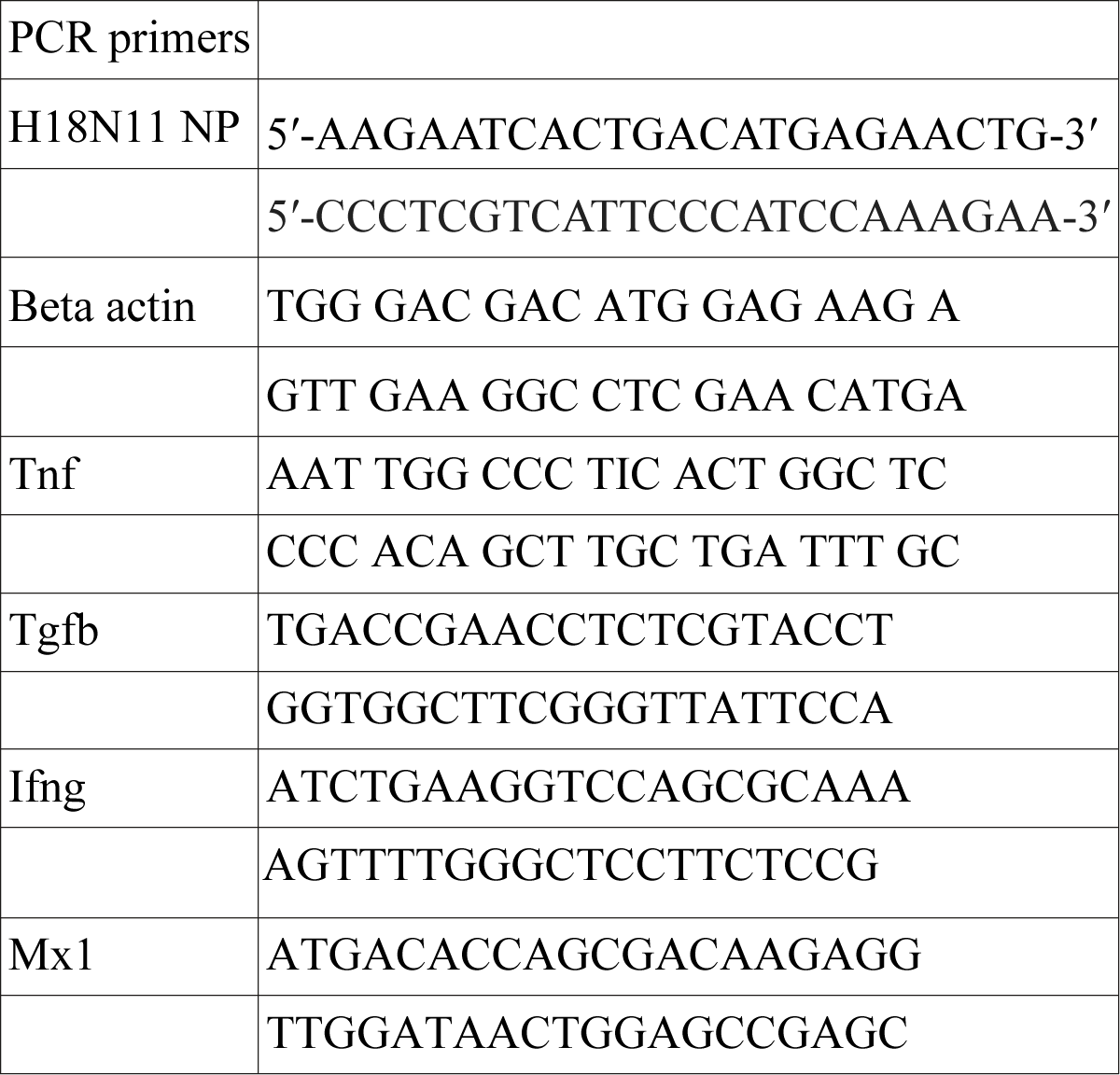

### Metabolomics

Plasma (EDTA, Greiner BioOne), rectal swabs (Puritan 5.5” Sterline Mini-tip Rayon swabs, #25-800-R50), and liver were collected for metabolomics analysis from all bats. Plasma and rectal samples collected during regular sampling were immediately placed into cold methanol then frozen until analysis. Liver sections were collected during necropsy. Time from sample collection until immersion in cold methanol was standardized.

For all metabolite extraction methods, derivatization, and LCMS methods, LCMS grade solvents were used. Tributylamine and all synthetic molecular references were purchased from Millipore Sigma. LCMS grade water, methanol, isopropanol and acetic acid were purchased through Fisher Scientific.

Plasma samples in methanol were brought to 800 µL with 1:1 methanol:water and 400 µL of chloroform was added. Liver samples were bead beaten in 400 µL methanol. Once samples were homogenized, 400 µL of water and 400 µL of chloroform were added. Rectal swabs were collected in 500 µL of methanol to which 500 µL of water and 500 µL of chloroform were added. All samples were shaken 30 minutes at 4 °C and centrifuged at 16000xg for 20 min. The top (aqueous) layer was taken for downstream analysis.

Short chain fatty acids (SCFA) were derivatized with O-benzylhydroxylamine (O-BHA) to prevent vaporization using previously established methods ^37^. Briefly, to 35 µL of each extracted liver and rectal swab sample was added 10 µL of 1M O-BHA and 10 µL of 1M 1-Ethyl-3-(3-dimethylaminopropyl)carbodiimide both in a reaction buffer consisting of freshly prepared 1M pyridine and 0.5 M hydrochloric acid in water. Samples were shaken at room temperature for 2 hours and then quenched via addition of 50 µL 0.1 % formic acid. Derivatized carboxylic acids including SCFA’s were extracted by adding 400 µL ethyl acetate to each reaction. Following mixing and centrifugation to induce layering, the upper (organic) layer was collected and dried under vacuum. Samples were resuspended in 300 µL of water for LCMS injection.

All samples were separated using a Sciex ExionLC™ AC system and measured using a Sciex 5500 QTRAP® mass spectrometer. Aqueous-soluble metabolites were analyzed using an ion pairing method^38^. Quality control samples consisting of a mixture of ten or more experimental samples from each biological matrix were injected regularly to control for instrument stability. Samples were separated on a Waters™ Atlantis T3 column (100Å, 3 µm, 3 mm X 100 mm) using a binary gradient from 5 mM tributylamine, 5 mM acetic acid in 2% isopropanol, 5% methanol, 93% water (v/v) to 100% isopropanol over 15 minutes. Two distinct MRM pairs in negative mode were used to measure each metabolite.

Derivatized SCFA samples were separated on a Waters™ Atlantis dC18 column (100Å, 3 µm, 3 mm X 100 mm) and eluted using a 6 min gradient of 5-80 % B with buffer A as 0.1 % formic acid in water and B as 0.1 % formic acid in methanol. SCFA and central metabolic carboxylic acids were detected using MRMs from previously established methods and identity was confirmed by comparison to derivatized standards including propionate, valerate, isovalerate, butyrate, isobutyrate, and acetate ^37,38^.

All peaks were picked and integrated using MultiQuant® Software 3.0.3. Signals were filtered by removing signals with greater than 50% missing values and the remaining missing values were replaced with the lowest registered signal value. All datasets were corrected for coherent QC variance via a linear nearest neighbors method. Metabolites with multiple MRMs were quantified with the higher signal to noise signal. Filtered datasets were normalized using the total signal sum prior to analysis. SCFA datasets were stitched to their corresponding polar metabolite dataset via common signals for malate in both methods.

### Antioxidant enzymes

We measured the activity of superoxide dismutase (SOD; Cayman, item no. 706002) and glutathione peroxidase (GPx; Cayman, item no. 703102) in red blood cell lysate following manufacturer’s instructions. 400uL of ice cold Ultrapure water (Cayman item #400000) was added to red blood cell lysate for 1 minute, briefly vortexed, centrifuged, and supernatant removed to use for each assay. Total protein in lysate was quantified using a BCA kit (Pierce BCA Protein Assay Kit #23225). SOD and GPx were both standardized to total protein.

### Data Analysis

Statistics were done using R 4.3.1 (R Core Team 2023-06-16) and the nlme (v 3.1-162, ^39^) and mixOmics ^40^. For metabolomics data, we also used MarkerView® Software 1.3.1 and MetaboAnalyst 5.0 ^41^.

All data were assessed for normality and either transformed or analyzed using non-parametric tests. Unless noted otherwise, diet-specific differences were compared using ANOVAs follow by Tukey HSD if significant or Kruskal-Wallis followed by pairwise Wilcox test if data were not normal based on Shapiro-Wilk test. Between day differences (within a given diet) were tested using paired t-tests or Mann-Whitney test if data were not normal based on Shapiro-Wilk test. P-values were corrected for multiple tests when multiple tests were run (FDR correction).

Data on weight were assessed using linear mixed effects models (“lme”, nlme) to determine if weight changed from the diet (day – 21 to day 0) or due to infection (day 0 to day 20). Gene expression data were normalized as log fold change relative to pre-infection values of all bats. For calculation of proportion of food consumed, pre-infection we did not weigh all food prior to giving it to the bats. In order to calculate an estimate of the proportion consumed, we used the average weight of food provided post-infection to the bats. Post-infection, food was weighed before and after so the actual proportion consumed was calculated.

## Supplementary Figures

**Supplementary Figure 1:**
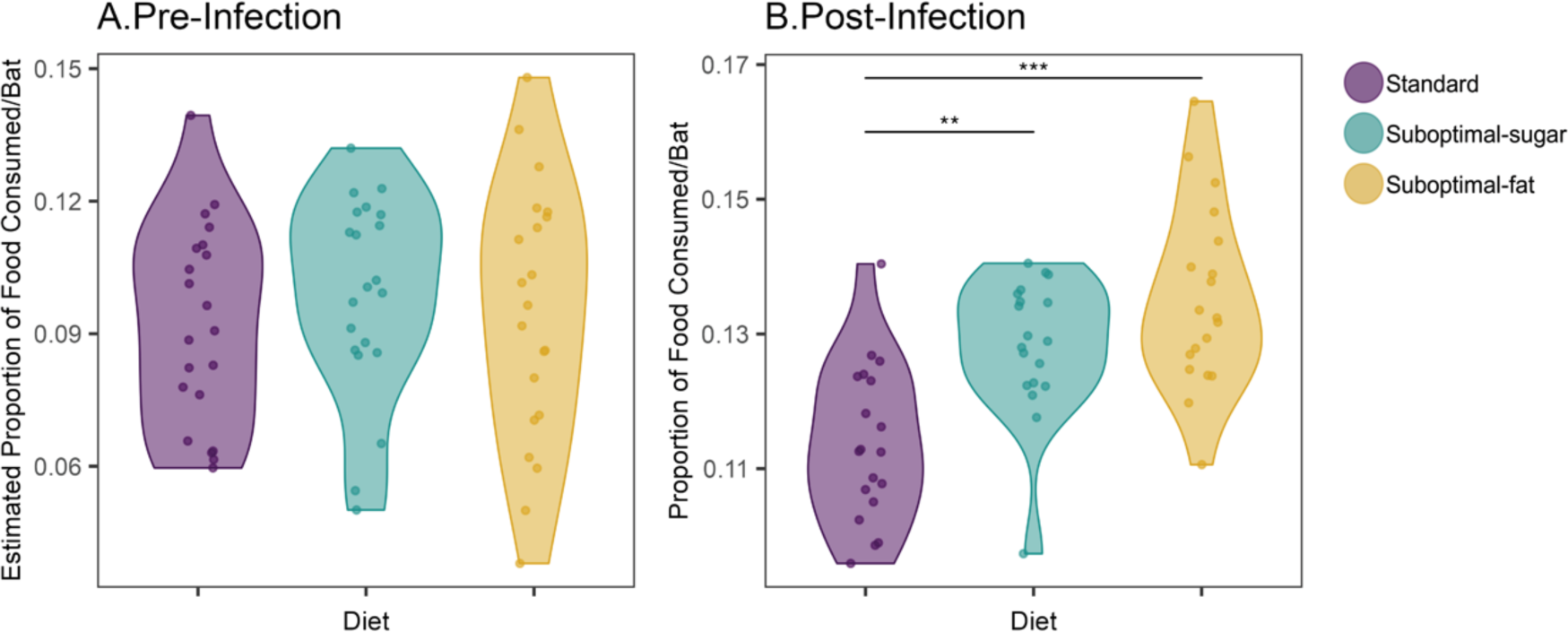
Food consumed per cage, normalized to number of bats. Food consumed, normalized by number of bats per cage instead of by average weight. (A) Pre-infection estimated grams of food consumed per bat, based on weight of leftover food. (B) Post-infection grams of food consumed per bat, based on difference between weight of food given and weight of food leftover. Both suboptimal diet cages consumed more food overall than standard diet bats (ANOVA, F = 16.15, df = 2, p < 0.001). *Asterisks indicate:* p < 0.0005 = ***, p < 0.005 = **, p < 0.05 = *, p < 0.1 =.

**Supplemental Figure 2.**
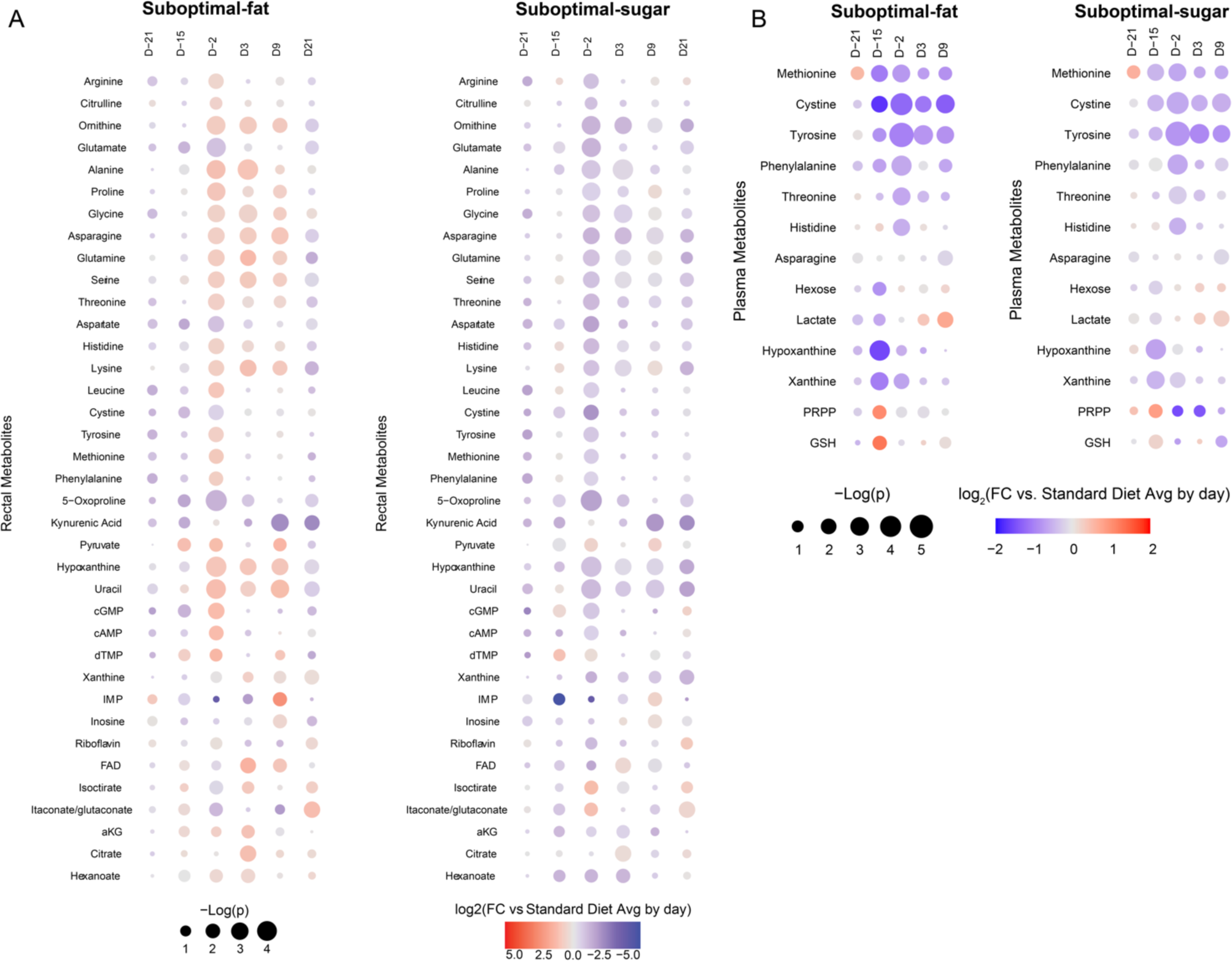
Diet differences in metabolite patterns stable by infection and maintained post-infection. Rectal (A, left) and plasma (B, right) metabolites throughout experiment that were significantly different between diets (based on passed 10% FDR filter). Color indicates log2-fold change relative to standard diet bats on each day and size of dot indicates -log(p value).

**Supplemental Figure 3:**
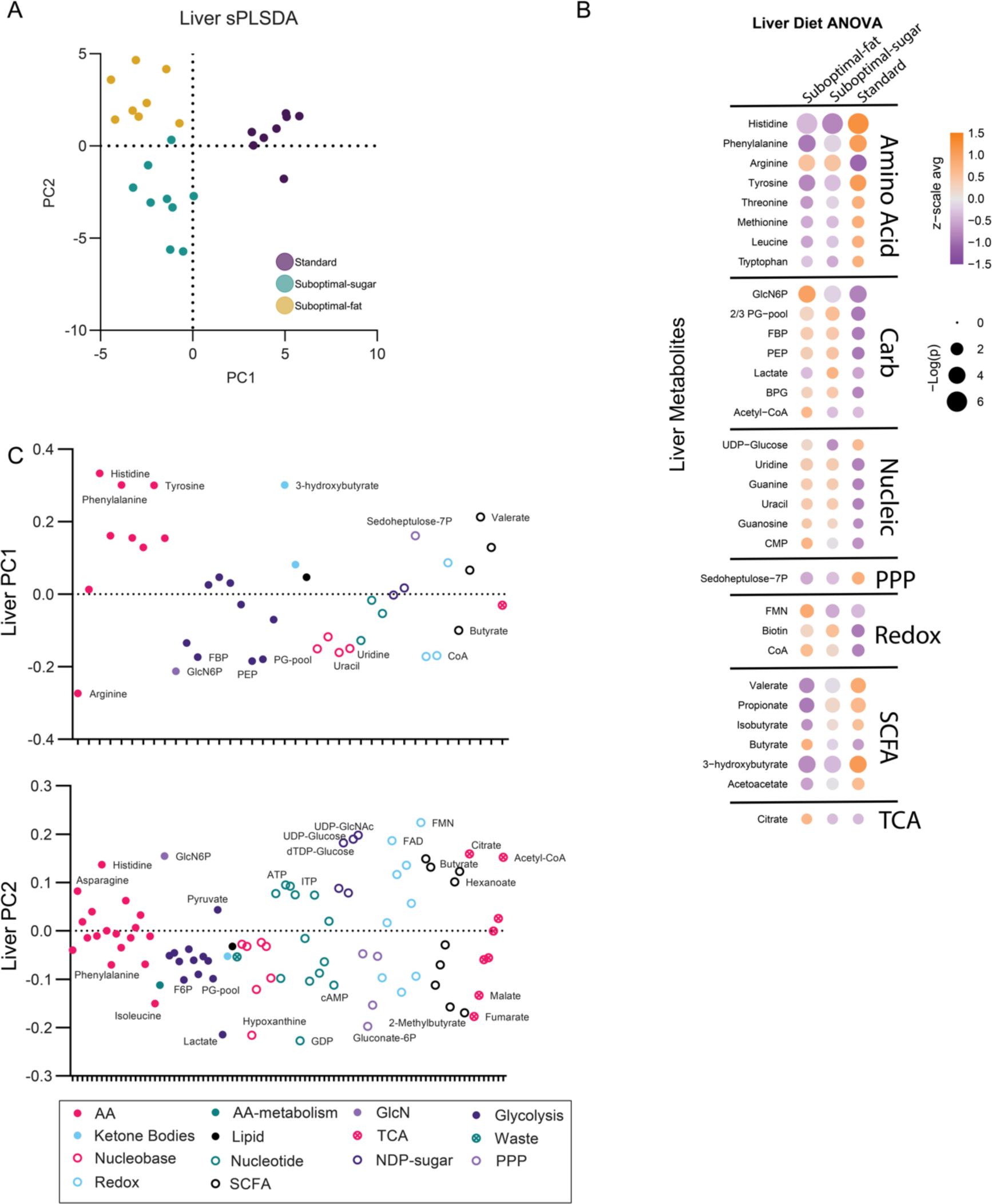
Liver metabolites reflect main energetic source of diet. (A) Sparse partial least squares regression (sPLSDA) classification based on liver samples from bats in each diet. Each point represents an individual bat. (B) Metabolites that were significantly different between diets (based on 10% FDR filter). Color indicates z-scale average and size of dot indicates -log(p value). (C) Loadings of metabolites from first two axes of sPLSDA plot (PC1 and PC2).

## Notes

### Competing Interest Statement

The authors have declared no competing interest.

## References

1. Gottdenker, N. L., Streicker, D. G., Faust, C. L. & Carroll, C. R. Anthropogenic Land Use Change and Infectious Diseases: A Review of the Evidence. Ecohealth 11, 619–632 (2014).

2. Allen, T. et al. Global hotspots and correlates of emerging zoonotic diseases. Nat. Commun. 8, 1–10 (2017).

3. Jones, B. A. et al. Zoonosis emergence linked to agricultural intensification and environmental change. Proc. Natl. Acad. Sci. U. S. A. 110, 8399–8404 (2013).

4. Plowright, R. K. et al. Land use-induced spillover: a call to action to safeguard environmental, animal, and human health. Lancet Planet. Heal. 5, e237–e245 (2021).

5. Demas, G. & Nelson, R. Ecoimmunology. (Oxford University Press, 2012).

6. Lochmiller, R. L. & Deerenberg, C. Trade-offs in evolutionary immunology: Just what is the cost of immunity? Oikos 88, 87–98 (2000).

7. Schaible, U. E. & Kaufmann, S. H. E. Malnutrition and infection: Complex mechanisms and global impacts. PLoS Med. 4, 0806–0812 (2007).

8. Kumar, A. et al. Impact of nutrition and rotavirus infection on the infant gut microbiota in a humanized pig model. BMC Gastroenterol. 18, 1–17 (2018).

9. Maier, E. A. et al. Protein-energy malnutrition alters IgA responses to rotavirus vaccination and infection but does not impair vaccine efficacy in mice. Vaccine 32, 48–53 (2013).

10. Taylor, A. K. et al. Protein energy malnutrition decreases immunity and increases susceptibility to influenza infection in mice. J. Infect. Dis. 207, 501–510 (2013).

11. Port, J. R. et al. High-fat high-sugar diet-induced changes in the lipid metabolism are associated with mildly increased covid-19 severity and delayed recovery in the syrian hamster. Viruses 13, 2506 (2021).

12. Eby, P. et al. Pathogen spillover driven by rapid changes in bat ecology. Nature 613, 340–344 (2023).

13. McKee, C. D. et al. Nipah Virus Detection at Bat Roosts after Spillover Events, Bangladesh, 2012-2019. Emerg. Infect. Dis. 28, 1384–1392 (2022).

14. Wood, M. R., de Vries, J. L., Epstein, J. H. & Markotter, W. Variations in small-scale movements of Rousettus aegyptiacus, a Marburg virus reservoir across a seasonal gradient. Front. Zool. 20, 1–17 (2023).

15. Markus, N. & Hall, L. Foraging behaviour of the black flying-fox (Pteropus alecto) in the urban landscape of Brisbane, Queensland. Wildl. Res. 31, 345–355 (2004).

16. Field, H. E. et al. Landscape Utilisation, Animal Behaviour and Hendra Virus Risk. Ecohealth 13, 26–38 (2016).

17. Parry-Jones, K. A. & Augee, M. L. The diet of flying-foxes in the Sydney and Gosford areas of New South Wales, based on sighting reports 1986-1990. Aust. Zool. 27, 49–54 (1991).

18. Manning, R. Pollen Analysis of Eucalypts in Western Australia. http://www.rirdc.gov.au (2001).

19. Gosper, C. R. & Vivian-Smith, G. Fruit traits of vertebrate-dispersed alien plants: Smaller seeds and more pulp sugar than indigenous species. Biol. Invasions 12, 2153–2163 (2010).

20. Lescano, C. H. et al. Nutritional and chemical characterizations of fruits obtained from Syagrus romanzoffiana, Attalea dubia, Attalea phalerata and mauritia flexuosa. J. Food Meas. Charact. 12, 1284– 1294 (2018).

21. Sadler, A. J. & Williams, B. R. G. Interferon-inducible antiviral effectors. Nat. Rev. Immunol. 8, 559–568 (2008).

22. Cross, P. C., Lloyd-Smith, J. O., Johnson, P. L. F. & Getz, W. M. Duelling timescales of host movement and disease recovery determine invasion of disease in structured populations. Ecology Letters vol. 8 587– 595 (2005).

23. Reiter, J. L., Crissey, S. D. & Brewer, B. A. Effect of Dietary Protein Level on Dry Matter Intake and Body Mass of. in American Association of Zoo Veterinarians 333–339 (1994).

24. Plowright, R. K. et al. Pathways to zoonotic spillover. Nat. Rev. Microbiol. 15, 502–510 (2017).

25. Farré, R., Fiorani, M., Rahiman, S. A. & Matteoli, G. Intestinal permeability, inflammation and the role of nutrients. Nutrients 12, 1–18 (2020).

26. Lee, J. S. et al. Hypoxanthine is a checkpoint stress metabolite in colonic epithelial energy modulation and barrier function. J. Biol. Chem. 293, 6039–6051 (2018).

27. Zoghroban, H. S., Ibrahim, F. Mk., Nasef, N. A. & Saad, A. E. The impact of L-citrulline on murine intestinal cell integrity, immune response, and arginine metabolism in the face of Giardia lamblia infection. Acta Trop. 237, 106748 (2023).

28. Trompette, A. et al. Dietary Fiber Confers Protection against Flu by Shaping Ly6c− Patrolling Monocyte Hematopoiesis and CD8+ T Cell Metabolism. Immunity 48, 992–1005.e8 (2018).

29. Guito, J. C., Prescott, J. B., Arnold, C. E., Palacios, G. F. & Towner, J. S. Asymptomatic Infection of Marburg Virus Reservoir Bats Is Explained by a Strategy of Immunoprotective Disease Tolerance Asymptomatic Infection of Marburg Virus Reservoir Bats Is Explained by a Strategy of Immunoprotective Disease Tolerance. Curr. Biol. 1–14 (2021) doi:10.1016/j.cub.2020.10.015.

30. Mordstein, M. et al. Interferon-λ contributes to innate immunity of mice against influenza A virus but not against hepatotropic viruses. PLoS Pathog. 4, (2008).

31. Johnson, A. M. F. et al. High fat diet causes depletion of intestinal eosinophils associated with intestinal permeability. PLoS One 10, e0122195 (2015).

32. Serino, M. et al. Metabolic adaptation to a high-fat diet is associated with a change in the gut microbiota. Gut 61, 543–553 (2012).

33. Rohr, M. W., Narasimhulu, C. A., Rudeski-Rohr, T. A. & Parthasarathy, S. Negative Effects of a High-Fat Diet on Intestinal Permeability: A Review. Adv. Nutr. 11, 77–91 (2020).

34. Agans, R. et al. Dietary fatty acids sustain the growth of the human gut microbiota. Appl. Environ. Microbiol. 84, (2018).

35. Ciminski, K. et al. Bat influenza viruses transmit among bats but are poorly adapted to non-bat species. Nat. Microbiol. 4, 2298–2309 (2019).

36. Zhou, B. et al. Characterization of Uncultivable Bat Influenza Virus Using a Replicative Synthetic Virus. PLoS Pathog. 10, e1004420 (2014).

37. Zeng, M. & Cao, H. Fast quantification of short chain fatty acids and ketone bodies by liquid chromatography-tandem mass spectrometry after facile derivatization coupled with liquid-liquid extraction. J. Chromatogr. B Anal. Technol. Biomed. Life Sci. 1083, 137–145 (2018).

38. McCloskey, D., Gangoiti, J. A., Palsson, B. O. & Feist, A. M. A pH and solvent optimized reverse-phase ion-paring-LC–MS/MS method that leverages multiple scan-types for targeted absolute quantification of intracellular metabolites. Metabolomics 11, 1338–1350 (2015).

39. J, P., D, B. & R Core Team,. nlme: Linear and Nonlinear Mixed Effects Models. (2023).

40. Rohart, F., Gautier, B., Singh, A. & Lê Cao, K. A. mixOmics: An R package for ‘omics feature selection and multiple data integration. PLoS Comput. Biol. 13, (2017).

41. Pang, Z. et al. MetaboAnalyst 5.0: Narrowing the gap between raw spectra and functional insights. Nucleic Acids Res. 49, W388–W396 (2021).

